# BRAF-V600E-Mediated Erk Activation Promotes Sustained Cell Cycling and Broad Transcriptional Changes in Neonatal Cardiomyocytes

**DOI:** 10.1101/2022.02.28.482357

**Authors:** Nicholas Strash, Sophia DeLuca, Geovanni L. Janer Carattini, Yifan Chen, Jacob Scherba, Mehul Jain, Ramona Naseri, Tianyu Wu, Nenad Bursac

**Affiliations:** Department of Cell Biology, Duke University, Durham NC; Department of Biomedical Engineering, Duke University, Durham NC

**Keywords:** MAPK, ERK, engineered cardiac tissue, proliferation, hypertrophy, glycolysis

## Abstract

Mitogens capable of promoting cardiomyocyte proliferation represent important targets for functional heart regeneration following myocardial infarction. We previously described an ERK-dependent pro-proliferative tissue phenotype following overexpression of constitutively-active (ca) human ERBB2 in both neonatal rat ventricular myocytes (NRVMs) and human iPSC-derived cardiomyocytes (hiPSC-CMs). Since ERBB2 canonically regulates multiple other pathways in addition to ERK, it is unclear whether ERK activation alone can drive CM proliferation. Here, we activated ERK in a targeted fashion by CM-specific lentiviral expression of a constitutively active mutant of BRAF, BRAF-V600E (caBRAF), in cultured NRVMs and examined the effects on engineered NRVM tissue proliferation, morphology, and function. caBRAF expression induced ERK activation, tissue growth, loss of contractile function, and increased tissue stiffness, all of which were sustained for at least 4 weeks *in vitro*. From bulk RNA-sequencing analysis of engineered tissues, we found that caBRAF had broad transcriptomic effects on CMs and induced a shift to glycolytic metabolism. Together, this work shows that direct ERK activation is sufficient to modulate CM cycling and functional maturation in a cell-autonomous fashion and could offer a potential target for cardiac regenerative therapies.

## 1. Introduction

Myocardial infarction results in the permanent loss of CMs and decline in heart function. Promising strategies to replace the lost heart tissue and improve cardiac function include cell therapies to engraft functional CMs differentiated from PSCs^1-3^ or gene therapies to drive endogenous CM proliferation^4^. Understanding the mechanisms that regulate CM proliferation is of paramount importance for both therapeutic strategies as the recovery of cardiac muscle mass and function will be directly proportional to the numbers of exogenously engrafted CMs or CM number resulting from inducing endogenous proliferation.

The mitogen-activated protein kinase (MAPK) signaling pathway is a widely studied, complex signaling pathway which governs multiple biological processes essential for both embryonic development and maintenance of tissue homeostasis^5-8^. A plurality of human cancers possess at least one mutation in a component of the MAPK signaling cascade resulting in dysregulation of its primary canonical effectors, extracellular signal-regulated kinases (ERKs)^9, 10^. In healthy cells, ERK activity is initiated by upstream kinases, which are influenced by a variety of extracellular and intracellular signaling molecules. Constitutive activation of ERK in response to transient or sustained growth factor stimulation is prevented by inhibitory negative feedback that tightly regulate ERK activity^11, 12^. BRAF, a commonly-mutated protein in multiple cancer types, is a serine/threonine protein kinase within the canonical MAPK pathway that is responsible for regulating the kinase MEK which, in turn, activates ERK. The most common activating somatic mutation of BRAF is the V600E mutation which is known to evade negative feedback inhibition and as a result is highly tumorigenic^13^. Since BRAF is thought to only activate the MEK/ERK signaling axis, BRAF-V600E (caBRAF) can be used as means to study targeted activation of ERK^14^.

When activated in the heart, postnatally or *in vitro*, ERK has been paradoxically characterized as a promoter of both CM hypertrophy^7, 15-17^ and proliferation^18-21^. ERK dysregulation in the heart during development has been linked to congenital cardiac defects characterized by concentric hypertrophy, as reported for patients with Noonan syndrome, Costello syndrome, and cardio-facio-cutaneous syndrome^6^. One study utilized patient-derived hiPSC lines with germline BRAF-activating mutations (T599R and Q257R) to show that CMs differentiated from these lines displayed a phenotype reflective of hypertrophic cardiomyopathy^22^. In another study, engineered cardiac tissues (ECTs) generated from BRAF-T599R hiPSCs exhibited similar functional changes^23^.

In this report, we utilized a three-dimensional (3D) NRVM ECT culture system exhibiting advanced maturation and function^24-26^ to study the structural and functional effects of targeted ERK activation induced by CM-specific lentiviral expression of caBRAF. We observed pro-proliferative and anti-maturation effects on CMs that yielded profound tissue growth and functional deficit in ECTs lasting for at least 4 weeks in culture. RNA-sequencing analysis of control and caBRAF tissues revealed broad transcriptomic differences in cell metabolism and cell-matrix interactions that underlie the observed functional changes. Our *in vitro* studies suggest that sustained ERK activity can counter the natural maturation of postnatal CMs yielding a pro-growth phenotype of potential relevance for congenital heart diseases and future regenerative therapies.

## 2. Materials and Methods

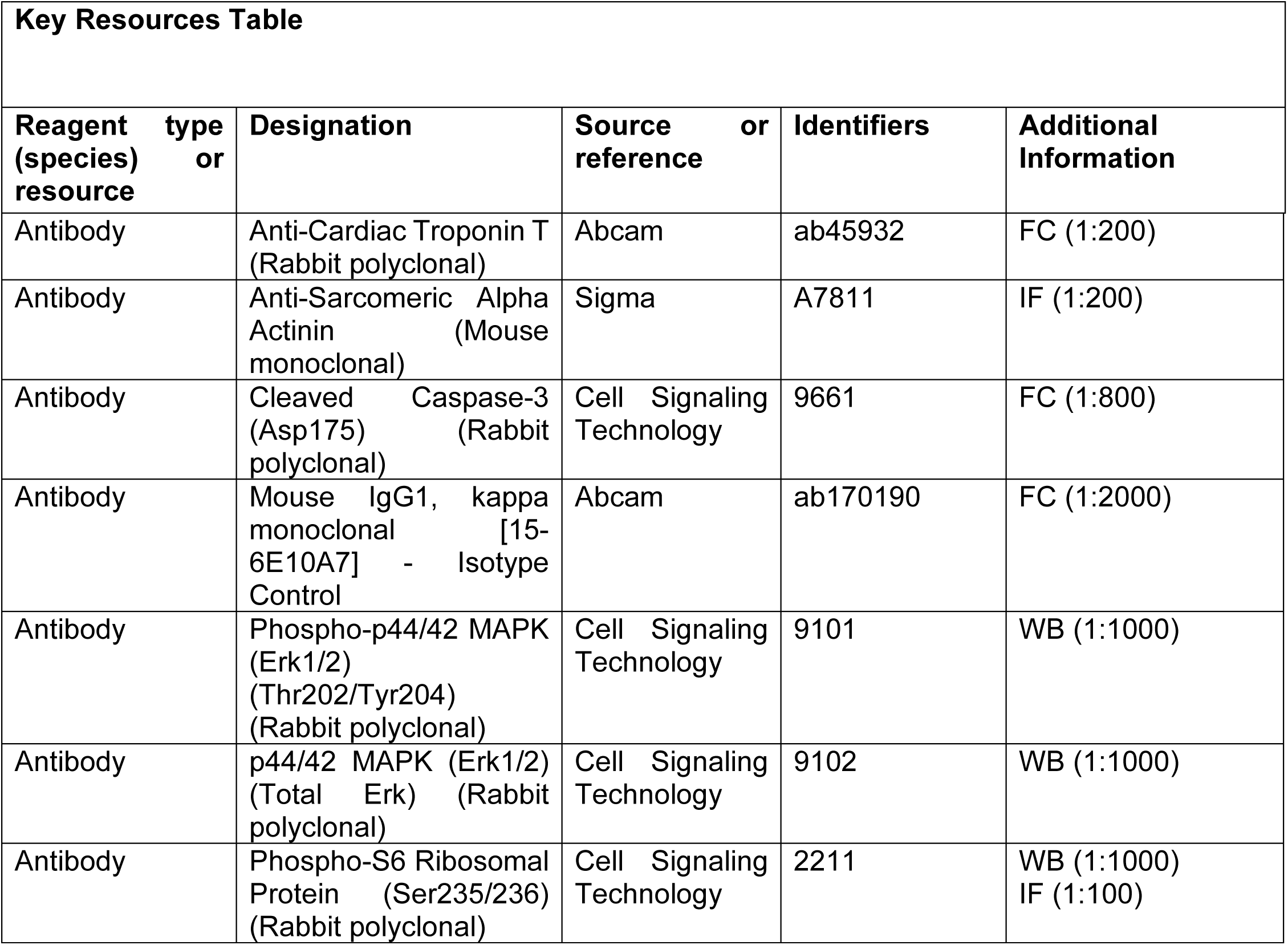
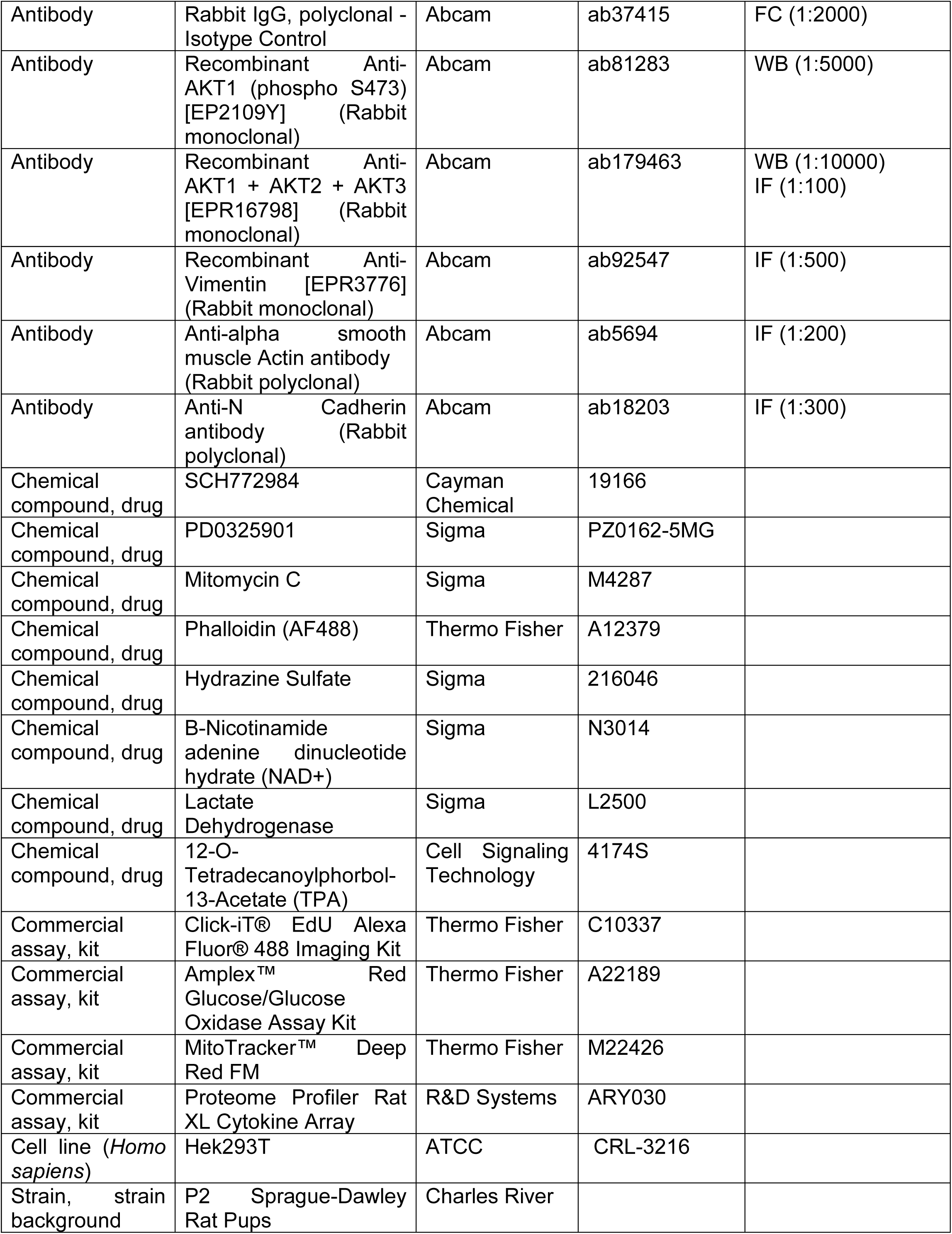
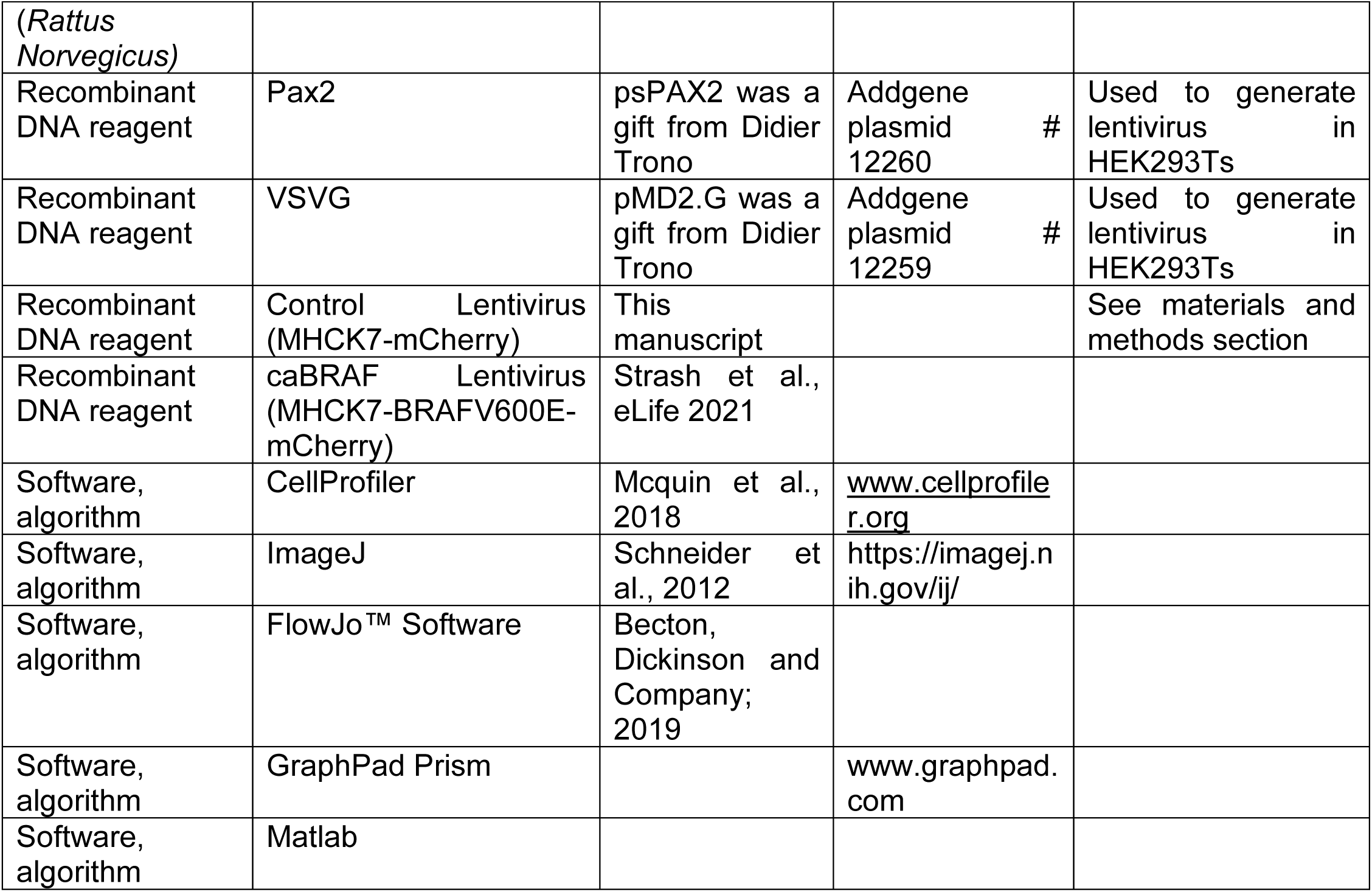

### NRVM Isolation and 2D culture

All animal procedures were performed in compliance with the Institutional Animal Care and Use Committee at Duke University and the NIH Guide for the Care and Use of Laboratory Animals. Neonatal rat ventricular myocytes (NRVMs) were isolated as previously described ^24, 25, 27^. Briefly, ventricles were harvested from P2 male and female Sprague-Dawley rat pups, minced finely, and pooled before overnight trypsin incubation at 4°C. The following day, the minced ventricular tissue was subjected to several collagenase digestion and filtering steps to yield single cell suspension. Cells were pre-plated for 1 hour to remove non-myocytes and enrich the NRVM population. The non-adherent cells were resuspended in 2D cardiac medium (DMEM, 10% FBS, penicillin (5 U/ml), vitamin B12 (2 µg/ml) and plated onto fibronectin-coated Aclar® coverslips at a density of 5×10^5^ cells per well of a 12-well plate. Twenty-four hours following plating the medium was changed to only include 5% FBS and full media changes were performed every other day.

### ECT Fabrication and 3D Culture

NRVM ECTs were prepared as previously described ^24-26^. Briefly, 6.5 × 10^5^ freshly isolated NRVMs were mixed with a fibrin-based hydrogel (2.5 mg/ml fibrinogen, 1 U/ml thrombin, 10% v/v Matrigel) and cast in PDMS tissue molds with two 2 mm x 7 mm troughs and a porous nylon frame. The molds containing the hydrogel-cell mixture were incubated at 37°C for 45 minutes to allow the hydrogel to fully polymerize and attach to the nylon frame. Tissues were then immersed in 3D cardiac medium (Low Glucose DMEM, 10% horse serum, 0.5% chick embryo extract, aminocaproic acid (1 mg/ml), ascorbic acid 2-phosphate sesquimagnesium salt hydrate (50 µg/ml), penicillin (5 U/ml), vitamin B12 (2 µg/ml)). The following day, the ECTs on frames were carefully removed from the molds and cultured in free-floating dynamic conditions on a rocker. Full media changes of 2 mL per well were performed every other day for 14 days. For Co-culture experiments, a non-transduced ECT was cultured with another ECT transduced with either Ctrl LV or caBRAF LV and the non-transduced ECT was analyzed.

### Migration assay in 2D NRVM cultures

For assaying cell migration, mitomycin C was added the day after seeding to inhibit fibroblast proliferation and ensure assessment of predominantly cardiomyocytes. The cells (1wk after seeding) were time-lapse imaged on a Dragonfly spinning disk confocal microscope (Andor), taking one image every 10 minutes for ∼7hr while maintaining CO_2_ and temperature.

### Small molecule Erki and MEki experiments

For experiments utilizing small molecule inhibitors of Erk or Mek, SCH772984 (Erki) or PD0325901 (Meki) were used. Both inhibitors were applied from day 8-14 after tissue generation at 100nM, and were compared to a DMSO vehicle control. For experiments using the small molecule 12-O-Tetradecanoylphorbol-13-acetate (TPA), the tissues were treated from day 2-14 after tissue generation with 10nM TPA and compared to a DMSO vehicle control. See Key Resources table for drug information.

### Cloning of Mitogen Constructs

For generation of the mitogen construct caBRAF, plasmid containing the gene sequence was used as PCR template for amplification of insert prior to cloning (see Key Resources Table for plasmids). PCR primers were designed to add complementary restriction site overhangs to the gene inserts which were also present in the MHCK7-MCS-P2A-mCherry backbone used for cloning the constructs. Standard restriction cloning was used to insert the gene fragments. Sanger sequencing was performed to ensure maintenance of reading frame and correct sequence.

### Preparation of Lentivirus Vectors

Lentiviral vectors were prepared as previously described ^28^. Briefly, Hek293T cells were cultured in high glucose DMEM containing 10% FBS and 1% Penicillin/Streptomycin. Plasmids (Construct plasmid, Pax2, and VSVG) were purified using midiprep before transfection into Hek293T cells at 65-75% confluence using Jetprime transfection reagent. Medium was changed 16 hours following transfection, and medium containing virus was harvested 3-4 days following initial transfection. Virus was purified by precipitation using 3 volumes medium to 1 volume 40% PEG-8000 at 4°C overnight, then pelleted by centrifugation at 1500xg for 45 minutes at 4°C. Precipitated virus was aliquoted and stored at −80°C before use. For NRVM monolayer experiments, viral suspension was added at the time of cell plating. For NRVM ECT experiments, lentiviral vectors were added to the hydrogel-cell mixture at the time of ECT fabrication to yield transduction efficiency between 60-80%.

### Flow Cytometry

NRVM monolayers were rinsed with PBS then dissociated using 0.05% Trypsin-EDTA at 37°C for 3 minutes, upon which monolayers were triturated several times to yield a single cell suspension. Trypsin was quenched with DMEM/F12 containing 20% FBS and 20µg/mL DNase I. The cell suspension was centrifuged at 300xg for 5 minutes, then resuspended in 4% PFA diluted in PBS. Cells were incubated in PFA for 10 minutes at RT, centrifuged again, then resuspended in PBS containing 5% FBS for storage at 4°C.

Cells were stained for flow cytometry after centrifugation at 300xg for 5 minutes to remove storage medium. If EdU staining was performed, cells were incubated with the EdU flow cytometry staining cocktail as per manufacturer protocol (ThermoFisher) and incubated in the dark for 30 minutes, then washed 2X by addition of PBS followed by centrifugation. Antibody staining was performed after EdU staining. For antibody staining, cells were resuspended in FACS buffer (PBS with 0.5% BSA, 0.1% Triton-X 100, 0.02% sodium azide). Primary antibodies including an isotype control were diluted in FACS buffer and incubated for 1 hour on ice. Cells were washed 2X with FACS buffer before addition of secondary antibodies and Hoechst diluted in FACS buffer. Secondary antibodies were incubated for 30 minutes at RT. Samples were run on a BD Fortessa X-20.

### Immunostaining and Imaging

Performed as previously described^29^, cell monolayers were fixed with 4% v/v PFA at room temperature for 15 minutes, then blocked in antibody buffer (5 w/v donkey serum, 0.1% v/v Triton X-100, in PBS) for 30 minutes at room temperature and incubated with primary antibodies for 30 minutes in antibody buffer. Primary antibody sources and dilutions are indicated in the Key Resources Table. The monolayers were washed with PBS before incubation with Alexa Fluor-conjugated secondary antibodies at 1:1000 and Hoechst at 1:200 in antibody buffer for 30 minutes. Monolayer samples were mounted using Fluoromount-G™ Mounting Medium and imaged using an Andor Dragonfly spinning disk confocal microscope.

Engineered ECTs were fixed with 2% v/v PFA on a rocking platform at 4°C overnight. For cross-sectional analysis, the fixed tissues were suspended in OCT and flash frozen in liquid nitrogen until solidified. The frozen tissue blocks were sectioned using a cryostat (Leica) into 10µm sections. ECT cross-sections were blocked in antibody buffer for two hours at room temperature. Whole bundles for longitudinal images were blocked overnight at 4°C. All samples were incubated with primary antibodies 4°C overnight in antibody buffer. Primary antibodies were used at the indicated dilutions in the Key Resources Table. Samples were incubated with Alexa Fluor-conjugated secondary antibodies at 1:1000 and Hoechst 1:200 in antibody buffer for 2.5 hours at room temperature for cross-sections and overnight at 4°C for whole bundles. Cross-sections and un-sectioned whole bundles were mounted with hard-set mounting medium (Antifade Glass) and imaged using an Andor Dragonfly spinning disk confocal microscope.

### qPCR

RNA was extracted using RNeasy Plus Mini Kit according to the manufacturer’s instructions (Qiagen). Total RNA was converted to cDNA using iScript cDNA synthesis kit (Bio-Rad). Standard qPCR reactions with 5 or 10 ng cDNA per reaction were performed with iTaq Universal SYBR Green Supermix (Bio-Rad) in the CFX Connect Real-Time PCR Detection System. All primers used are listed in qPCR Primer Table in Key Resources Table.

### RNA-sequencing

NRVM ECTs were pooled (6-8 per sample, each sample was from a separate NRVM isolation from a total of 3 isolations) and homogenized in RLT buffer (Qiagen) using green RINO homogenization tubes following the manufacturer’s recommendations. RNeasy Fibrous Tissue Mini Kit (Qiagen) was used to isolate RNA from the homogenized tissue, which was then sent for RNA-sequencing by Genewiz. Genewiz performed library preparation and sequencing; briefly, rRNA was removed using PolyA selection for mRNA species and sequencing was performed using Illumina HiSeq, 2×150bp paired-end reads with 20-30 million reads per sample. Read quality was confirmed using FastQC software^30^. Alignment to the *R. norvegicus* genome assembly was performed using the Rsubread package^31^. Differential expression was performed using DESeq2^32^. Significantly differentially expressed genes were classified as: absolute value of Log_2_FC > 1.5 and p_adj_ < 0.01.

### Western Blot

To isolate total protein from NRVM ECTs, cells were rinsed twice with ice cold PBS before lysis with RIPA buffer containing protease inhibitor cocktail (Sigma P8340) and phosphatase inhibitor cocktail 3 (Sigma P0044). Cells were incubated on ice for 10 minutes, then lysates were collected and spun down at 10,000xg to pellet debris. Supernatants were measured using BCA assay to determine total protein concentration. Thirty μg of each sample was run on a 4-12% gradient gel with Tris-Glycine-SDS running buffer at 100V for 1-1.5 hours depending on the size of proteins being separated. Proteins were transferred to 0.45µm PVDF membranes for Western blot at 4°C at 60V for 2 hours. Membranes were blocked overnight in 3% BSA in Tris-buffered saline (TBS). Membranes were cut such that multiple size proteins could be blotted from the same membranes. Primary antibodies were diluted in 3% BSA and incubated with membranes overnight at 4°C. Membranes were washed 3X with TBS containing 0.1% Tween-20 (TBS-T) before incubation with HRP-conjugated secondary antibodies. Membranes were washed 3X with TBS-T before incubation in SuperSignal™ West Pico PLUS Chemiluminescent Substrate for 5 minutes. Membranes were imaged with a Biorad ChemiDoc using signal accumulation mode for up to 2 minutes. If an additional protein of interest was similar in size to the housekeeping gene (LamB1) or other proteins of interest, membranes were stripped following exposure for 10 minutes using Restore™ PLUS Western Blot Stripping Buffer. Membranes were then reblocked and re-probed as indicated above. Antibodies and their dilutions can be found in the Key Resources table.

### Conditioned media assays

For the cytokine array, R&D systems’ Proteome Profiler Rat XL Cytokine Array was used based on manufacturer’s recommendations. Membranes were imaged with a Biorad ChemiDoc using signal accumulation mode for up to 2 minutes. To measure the concentration of glucose in the collected culture medium, the Amplex Red glucose assay kit from Invitrogen was used following manufacturer’s recommendations. To measure the concentration of lactate in the collected culture medium, we performed a plate reader assay as we have used previously^33^. Briefly, a lactate dehydrogenase-catalyzed reaction with lactate, NAD+, and hydrazine was used, and the formation of reduced NAD was followed spectrophotometrically at 340nm kinetically up to 60 minutes. The concentration of lactate in the medium was then calculated based on the linear range of a standard curve.

### Force measurements

ECT force generation was measured using a custom-made force measurement setup consisting of an optical force transducer and linear actuator as previously described ^25^. In 37°C Tyrode’s solution, the ECT was pinned to chamber at one end and a PDMS float connected to a linear actuator controlled by Labview software at the other end. Using platinum electrodes, a 90V biphasic electrical pulse was applied for 5 ms at 2 Hz rate to induce contractions. The force measurements were performed at the ends of 4% stretch steps lasting 45 seconds until 12% stretch was reached. Stretch distance to achieve 4% stretch steps was determined based on the 7mm resting length for control tissues. Due to high tissue stiffness with caBRAF, 0% stretch was measured starting at the point of zero passive force, which was normally less than 7mm. Maximum twitch amplitude (occurring anywhere between 0% and 12% stretch), passive force-length curves, and parameters of twitch kinetics were derived as previously described using custom Matlab software ^34^.

## QUANTIFICATION AND STATISTICAL ANALYSIS

Statistical analysis was performed with GraphPad Prism software. Outliers were identified and removed using GraphPad Prism 8.3.0 ROUT method (Q=1%). Normality testing was done using the Shapiro-Wilk test and testing for equal variances was done using the Brown-Forsythe test. If data was not normally distributed, we performed logarithmic transformations and re-tested for normality and equal variances prior to performing the appropriate statistical test. All experiments were carried out in multiple cell batches. n’s are defined as being a single engineered tissue or single well in a culture dish.

### Image Analysis

Image analysis was performed using custom FIJI^35^ macros. Colocalization analysis between vimentin signal and nuclei as well as EdU signal and nuclei was performed to exclude proliferative fibroblasts from cardiomyocyte EdU quantification. A custom FIJI macro using auto-thresholding methods was used to determine ECT F-actin^+^ area and total cross-sectional area (CSA).

### Migration Analysis

Cell migration analysis on time lapse images was performed using the TrackObjects module in CellProfiler^36^ to determine cell position, RMS displacement, and cell speed. The resulting cell speed and position data was fit to a Persistent Random Walk model as shown previously^37, 38^ using custom Matlab code to calculate mean square displacement (MSD), persistence time and mean free path (MFP). For MSD analysis, cells tracked for less than 100 consecutive seconds were excluded from analysis. Migratory cells were considered to have a MFP > 0.1µm based on measurements of visually non-migratory cells.

## 3. Results

### 3.1 caBRAF expression promotes cell cycling, morphological changes, and functional deficit in NRVM ECTs

To ensure CM-specific transgene expression, we generated lentiviruses (LVs) in which the muscle-specific MHCK7 promoter^39^ drove expression of mCherry (control, Ctrl) or caBRAF-2A-mCherryNLS (Fig. 1-S1A). We first examined the effect of LV-expressed caBRAF in NRVM monolayers and found that sarcomere organization in CMs was disrupted by 1wk and further deteriorated by 2wks of culture (Fig. 1A). In a 3D NRVM ECT system, CM-specific expression of caBRAF was associated with larger tissue cross sectional area (CSA) by 1wk (Fig. 1B,C) and this morphological change persisted at 2wks of culture at which point formation of central acellular core was apparent (Fig. 1E,F). As previously shown^25, 40^, in Ctrl ECTs, aligned cardiomyocytes strongly expressing F-actin, but not vimentin, resided in the tissue interior and were surrounded by an outer layer of vimentin+ fibroblasts (Fig. 1B,E). caBRAF transduction yielded the occurrence of vimentin+ staining throughout the interior of the tissue, suggesting that caBRAF induced ectopic expression of vimentin in cardiomyocytes, consistent with the direct link between ERK activation and vimentin transcription shown in breast carcinoma^41^. To determine whether caBRAF also drives increased cell cycle activation in NRVM ECTs, we delivered a pulse of 10μM EdU for 48hr prior to tissue fixation and observed no difference in the rates of DNA synthesis at 1wk (Fig. 1D), but increased total and CM-specific EdU incorporation at 2wks of culture (Fig. 1G). Together, caBRAF expression led to rapid morphological changes in NRVMs associated with increased intermediate filament production and cell cycle activation.

**Figure 1.**
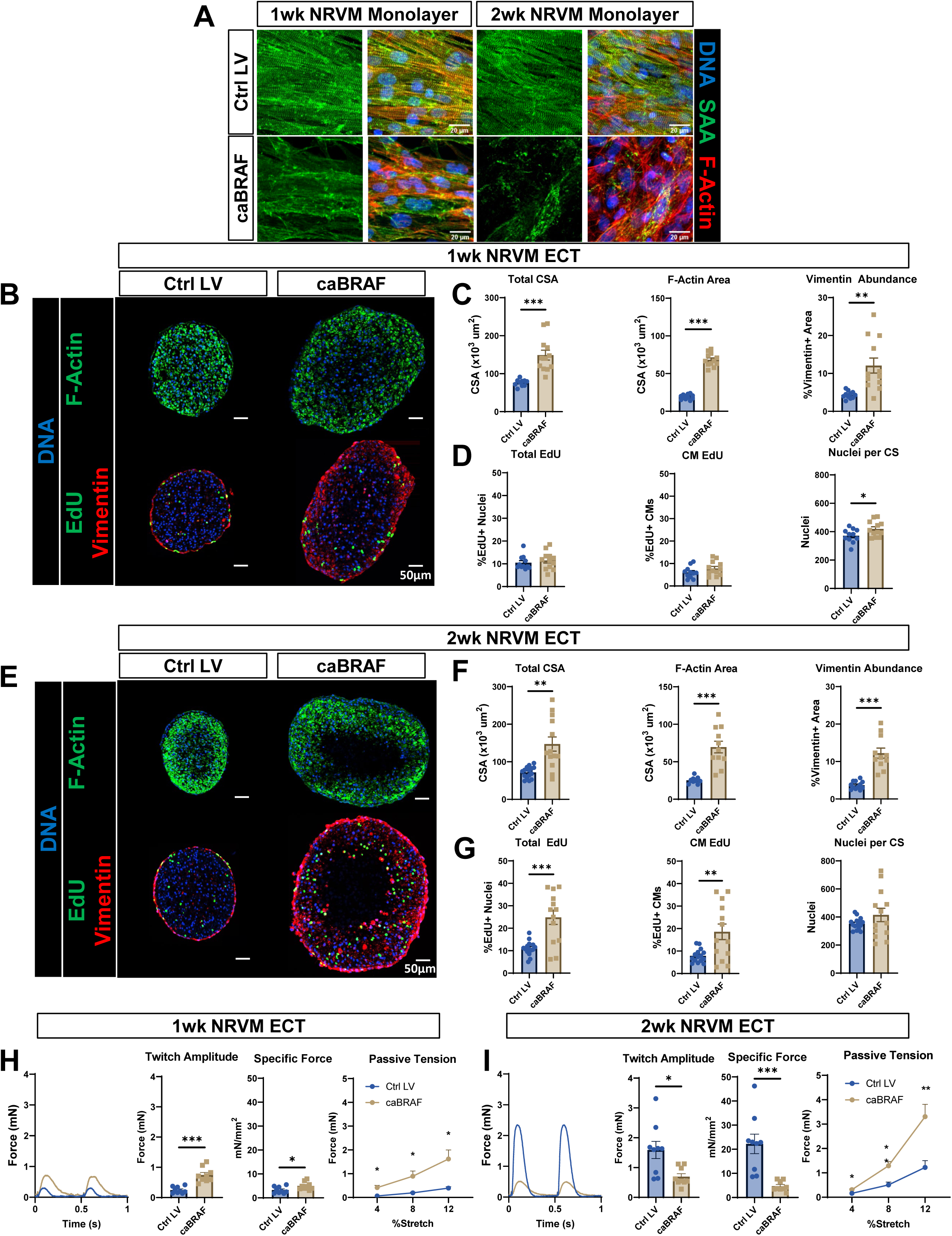
CaBraf Expression Induces Profound Morphological, Cel Cycling, and Functional Changes in NRVM ECTs with Time of Culture. (A) Representative images of (left) 1wk and (right) 2wk NRVM monolayers showing the deteriorating effects of caBRAF expression on sarcomere structure. Ctrl LV, control lentivirus; SAA, sarcomeric α-actinin. (B-G) Representative images of NRVM ECT cross-sections and corresponding morphological and cell cycling quantifications after 1 wk (B-D) and 2 wk (E-G) of culture. ECT, engineered cardiac tissue; CSA, cross-sectional area. (H-I) Representative twitch force traces at 2 Hz stimulation, twitch amplitudes, specific forces (force per CSA), and passive tension-length relationships in ECTs after 1 wk (H) and 2 wk (I) of culture. %Stretch values are shown relative to the initial testing length. Data: n=10-12 (C-D), n=11-15 (F-G), n=9 (H-I). Column graphs showing individual data points, mean ± SEM; Line plots, mean + SEM. *p < 0.05, **p < 0.01, ***p < 0.001 vs. Ctrl LV.

To explore functional consequences of caBRAF expression, we measured contractile function of NRVM ECTs using a custom force measurement system^24, 25, 42, 43^. After 1wk of culture, caBRAF ECTs showed increased contractile force (twitch) and specific force (force per CSA) amplitudes compared to control tissues (Fig. 1H). However, by 2wks of culture, caBRAF ECTs displayed significant decreases in both absolute (∼2.3-fold) and specific (∼4.7-fold) force compared to Ctrl ECTs (Fig.1I), which normally increase in contractile strength over the 2-week culture period, indicative of functional maturation^24, 25^. As in monolayer studies, sarcomere structure in caBRAF-expressing ECTs was disrupted at both 1 and 2wks (Fig. 1-S1B), which likely contributed to the development of the contractile deficit with time of culture. Additionally, caBRAF ECTs exhibited significantly increased passive tension at both 1 and 2wks of culture (Fig. 1H,I), similarly to the ERK-dependent tissue stiffening observed in our previous study with caERBB2 overexpression^29^. Finally, the observed contraction deficit was accompanied by slower twitch kinetics evident from both increased twitch rise and decay times that already showed changes after 1wk (Fig. 1-S1C-D). Collectively, in addition to profound morphological changes, caBRAF expression induced significant functional deficits in NRVM ECTs that developed by 2wks of culture and were evident from the decrease in contraction magnitude and speed of twitch kinetics and increase in tissue stiffness.

### 3.2 Phenotypic changes induced by caBRAF expression in NRVM ECTs persist and increase during prolonged culture

We then determined whether the caBRAF-induced phenotype at 1wk and 2wks of culture persisted longer-term. We found that compared to control ECTs, all morphological indices including F-actin^+^ CSA, total CSA, and vimentin expression remained increased in caBRAF at 3 and 4wk of culture (Fig. 1-S2AB,D,E). Although the EdU incorporation appeared to decrease over time compared to the earlier timepoints, caBRAF ECTs retained significantly higher rates of cell cycling at both 3 and 4wk (Fig. 1-S2C,F), with nuclei per CS progressively increasing over time. Additionally, caBRAF tissues showed persistent sarcomere disassembly at 3 and 4wk, with increased smooth muscle actin (SMA) expression observed at later time points of culture (Figure 1-S3A). We also found that contractile function of ECTs remained negatively impacted by caBRAF expression, including the persistent increase in the tissue tension (Fig. 1-S2G,H). Interestingly, twitch kinetics of caBRAF ECTs continued to deviate farther from Ctrl ECTs at the later time points, with caBRAF ECTs showing increased twitch duration from 288±5ms at 2wk to 374±22ms at 4wk, vs. Ctrl ECTs that exhibited stable twitch duration (189±3ms and 175±10ms at 2 and 4wks, respectively) (Fig. 1-S3B,C). Therefore, profound structural and morphological differences between caBRAF and Ctrl NRVM ECTs were maintained and even increased (for nuclei number, twitch decay time, and passive tension) with prolonged time of culture (Fig. 1-S4).

### 3.3 Bulk RNA-sequencing reveals profound transcriptomic changes in CMs caused by caBRAF expression

To gain further insights in molecular changes induced by caBRAF expression, we performed bulk RNA-sequencing on caBRAF and Ctrl ECTs at 1 and 2 wks of culture (Fig. 2). Broad differential clustering of expressed genes between Ctrl and CaBRAF tissues was apparent in heatmaps at both culture times (Fig. 2A). Specifically, 2664 and 2368 genes were differentially expressed between caBRAF and Ctrl ECTs in 1 and 2wk samples, respectively (Fig. 2B). Performing gene ontology (GO) analysis on these differentially expressed genes revealed that as expected, the GO terms MAPK Signaling Pathway and Cell Population Proliferation were upregulated, as well as Epithelial-Mesenchymal Transition (EMT) and ECM-Receptor Interactions, including multiple matrix metalloproteinases (Mmps) and integrins (Fig. 2-S1A). On the other hand, genes involved in Cardiac Muscle Development, Oxidative Phosphorylation, and Heart Contraction were downregulated. We also found predominant downregulation of major ion channels involved in CM electrical activity and calcium handling including SERCA2, RYR, and PLN (Fig. 2-S1B). Principal component analysis (PCA) showed that a majority of the variance between samples occurred due to caBRAF expression rather than length of culture (Fig. 2C).

**Figure 2.**
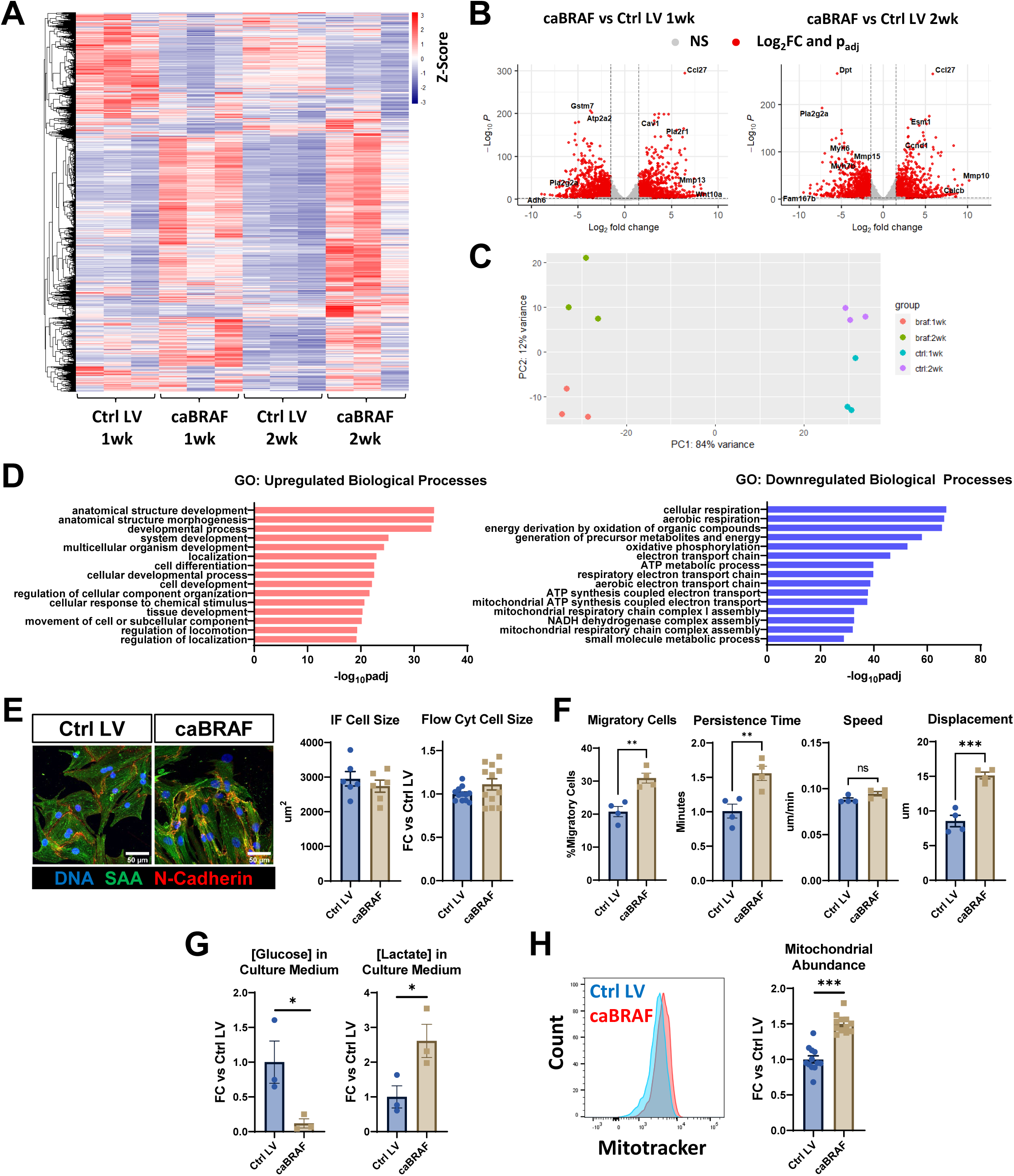
Transcriptomic Analysis of NRVM ECTs Reveals Effects of caBRAF Expression on CM Migration and Metabolism. (A-D) RNA-sequencing analysis showing (A) z-scores and clustering of all differentially expressed genes across groups, (B) volcano plots showing cutoff points for classifying a gene as differentially expressed (absolute value of Log_2_FC > 1.5 and p_adj_ < 0.01), (C) PCA clustering analysis, and (D) gene ontology (GO) analysis of the top upregulated and downregulated biological processes (at 2 wk) in caBRAF vs. Ctrl ECTs. (E) Representative images of 1wk NRVM monolayers (left) used for quantification of CM size (right). (F) Quantified parameters from cell migration analysis performed using live-imaging of 1 wk NRVM monolayers. (G) Glucose and lactate concentrations quantified in culture media of 1wk NRVM ECTs. (H) Representative flow cytometry-assessed fluorescence intensities of 1 wk NRVMs stained with Mitotracker® and corresponding quantifications of relative staining intensities. Data: n=6-12 (E), n=4 (F), n=3 (G), n=12 (H). Column graphs showing individual data points, mean ± SEM. *p < 0.05, **p < 0.01, ***p < 0.001 vs. Ctrl LV.

Notably, GO analysis returned the same or very similar differentially expressed pathways at the 1 and 2wk timepoints, hence, we focused on the pathways revealed by analysis of the 2wk tissues. Within differential GO terms of highest significance, we identified many upregulated processes related to development and locomotion (Fig. 2D), while downregulated GO terms pointed to profound metabolic changes, including decreased aerobic metabolism and mitochondrial function in caBRAF compared to Ctrl tissues (Fig. 2D). We then performed ChEA3 analysis^44^, which revealed likely transcription factors that mediated observed transcriptomic changes between caBRAF and Ctrl ECTs (Fig. 2-S1C). Of note, transcription factors known to mediate cardiac mesodermal specification and early development such as MEOX1, TCF15, GATA4, NKX2.5, and TBX20 were identified as likely regulators of the upregulated genes (Fig. 2-S1C)^45^. Taken together and consistent with our structural and functional studies, the transcriptomic analysis suggested that caBRAF expression induced anti-maturation effects in NRVM ECTs, characterized by a switch to an early cardiac developmental program, enhanced cell proliferation, ECM remodeling, and decreased cardiac function.

An unexpected result of our transcriptomic analysis was the lack of an obvious cardiac hypertrophic signature, as ERK has been widely reported to mediate CM hypertrophy^6, 7, 17, 46, 47^. To further explore whether caBRAF expression induced CM hypertrophy, we measured cell size in sparse NRVM monolayer cultures by quantitative immunostaining and found no size differences between caBRAF and Ctrl CMs, which was further confirmed using flow cytometry on NRVMs cultured in confluent monolayers (Fig. 2E). To additionally validate transcriptomic results, we performed live time-lapse microscopy imaging to quantify cell migration in NRVM monolayers since locomotion was a significantly upregulated process in GO analysis (Fig. 2D). Quantification of nuclear motion in monolayers over 7h (Fig. 2S2A) revealed an increased proportion of migratory cells, cell persistence time, and displacement induced by caBRAF expression (Fig. 2F), which was further evident from consistently right-shifted histograms of cell migration parameters (Fig. 2-S2B-C).

Since GO analysis suggested that aerobic metabolism was the most downregulated process in caBRAF vs. Ctrl ECTs, we additionally examined metabolic fuel consumption of ECTs by measuring the glucose and lactate concentrations in culture media. In caBRAF ECT media, glucose concentration was decreased and lactate concentration was increased, suggesting a shift favoring glycolytic metabolism (Fig. 2G), similar to findings in mice with CM-specific caERBB2 expression^48^. Interestingly, flow cytometric analysis using Mitotracker dye showed increased mitochondrial abundance in caBRAF NRVMs (Fig. 2H), consistent with known roles of ERK signaling in mitochondrial fission/fragmentation^49, 50^ and potentially signifying a compensatory mechanism where less efficient/fragmented mitochondria and decreased aerobic metabolism were countered by increased mitochondrial biogenesis. Together, results from the RNA-sequencing analysis suggesting caBRAF-induced CM migratory behavior and a metabolic shift towards glycolysis, but no CM hypertrophy, were all confirmed in the described follow-up functional studies.

### 3.4 Paracrine signals from caBRAF-expressing CMs contribute slower contraction kinetics of NRVM ECTs but not other phenotypic changes induced by caBRAF expression

A previous study suggested TGFβ-dependent paracrine roles of non-myocytes expressing mutated BRAF in hiPSC-CM hypertrophy^22^. This prompted us to examine the potential roles of paracrine signaling in driving the phenotypic changes in NRVMs observed with caBRAF expression. We first assessed differentially secreted cytokines in caBRAF ECTs at 1wk of culture using a qualitative cytokine array and found significant increases in GDF-15 and Osteoprotegerin, associated with TGFβ and TNFα signaling, respectively, as well as WNT-associated cytokines CCN1 and CCN4 (Fig. 3A). To further assess potential paracrine and autocrine effects from caBRAF expression, we cultured a caBRAF or a Ctrl ECT with a non-transduced ECT in the same well and performed structural and functional assessments after 2 weeks of culture^51^. Compared to co-culture with Ctrl ECTs, the non-transduced ECTs co-cultured with caBRAF ECTs exhibited a slight increase in CSA without other morphological changes (Fig. 3B,C), as well as a slight decrease in % Edu^+^ CMs without a change in total nuclei number (Fig. 3D). Functionally, compared to Ctrl ECTs, soluble factors from caBRAF ECTs led to a slight increase in contractile force generation in non-transduced ECTs, but no increase in specific force amplitude (Fig. 3E) or tissue stiffness (Fig. 3F). Furthermore, co-culture with caBRAF ECTs slightly increased twitch duration and decay time in non-transduced ECTs (Fig. 3G). Together, these studies suggested that soluble paracrine and autocrine signals from caBRAF-expressing CMs minimally contributed to the observed phenotype of NRVM ECTs, which instead was directly caused by CM-autonomous changes in MAPK signaling.

**Figure 3.**
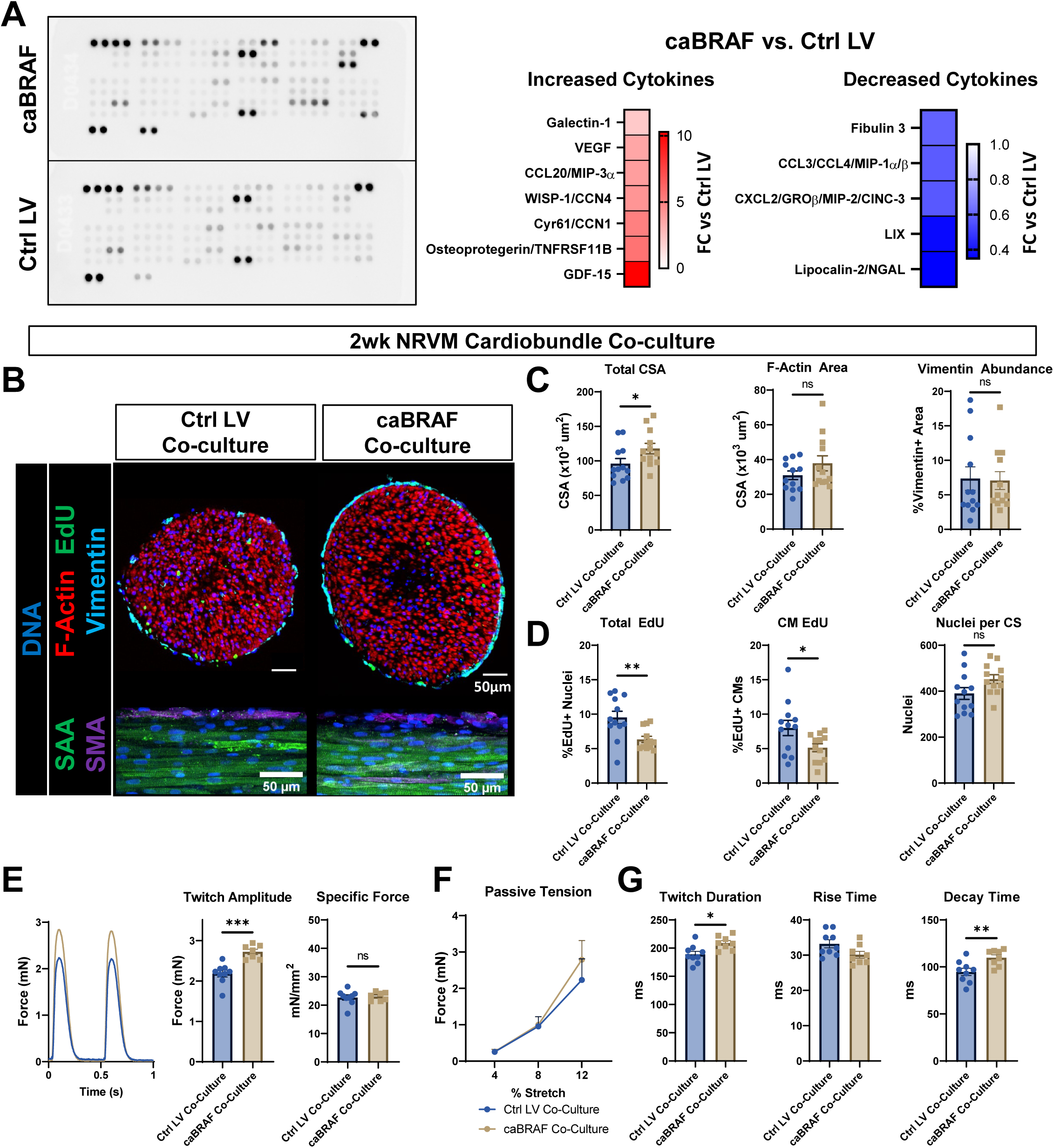
Secreted Factors from caBRAF ECTs are Minor Contributor to caBRAF Expression-induced Phenotype. (A) Image of a qualitative cytokine array used to analyze culture media from 1wk NRVM ECTs and corresponding quantifications of relative cytokine expressions in CaBRAF vs. Ctrl LV group. (B-D) Representative cross-section and whole-mount images (B) and corresponding quantifications of morphological (C) and cell cycling (D) parameters in non-transduced NRVM ECTs co-cultured for 2 wk with either Ctrl LV or caBRAF ECTs. SAA, sarcomeric α-actinin; SMA, smooth muscle actin; CSA, cross-sectional area. (E-G) Representative twitch force traces at 2 Hz stimulation and corresponding quantifications of twitch amplitudes and specific forces (E), passive tension-length relationships (F), and twitch kinetic parameters (G) in non-transduced NRVM ECTs co-cultured for 2 wk with either Ctrl LV or caBRAF ECTs. %Stretch values are shown relative to the initial testing length. Data: n=12 (C-D), n=7-9 (E-G). Column graphs showing individual data points, mean ± SEM. Line plots, mean + SEM. *p < 0.05, **p < 0.01, ***p < 0.001 vs. Ctrl LV.

### 3.5 Small molecule ERK activation recapitulates slower contraction kinetics of NRVM ECTs but not other phenotypic changes induced by caBRAF expression

To confirm that ERK signaling was activated by caBRAF expression, we performed Western blot analysis and found significant increases in total and phosphorylated (p-) ERK in both the 1wk and 2wk NRVM ECTs transduced with caBRAF, while AKT and p-AKT expression was not changed (Fig. 4A). We then aimed to determine whether small molecule activation of ERK would induce a similar phenotype in NRVM ECTs as observed with caBRAF expression. The small molecule 12-O-Tetradecanoylphorbol-13-acetate (TPA) has been frequently used to induce PKC-mediated, indirect ERK activation^52^. We treated the ECTs with 10nM TPA from culture day 2 (mimicking onset of lentivirally induced CaBRAF expression), every other day, until structural and functional assessments were performed at 2wk of culture. The TPA treatment induced moderate ERK activation based on Western blot analysis (Fig. 4-S1A,B), which was further confirmed from the increased expression of the ERK-dependent downstream genes *Etv4, Dusp6*, and *Spry4* (Fig. 4-S1C)^29, 53^. Expectedly, we also found increased *Fosb* and *Lgals3* expression, characteristic of activated PKC signaling (Fig. 4-S1C). However, despite the evidence for ERK activation, no changes in ECT morphology (Fig. 4C-D), cell cycling (Fig. 4E), contractile strength (Fig. 4E), or stiffness (Fig. 4F) akin to those induced by caBRAF expression were observed. On the other hand, the TPA treatment yielded an increase in both twitch duration and decay time (Fig. 4G). Collectively, these studies demonstrated that the caBRAF expression induced constitutive ERK activity in NRVM ECTs and that TPA-mediated ERK activation was unable to reproduce caBRAF effects aside from the slowing of twitch kinetics.

**Figure 4.**
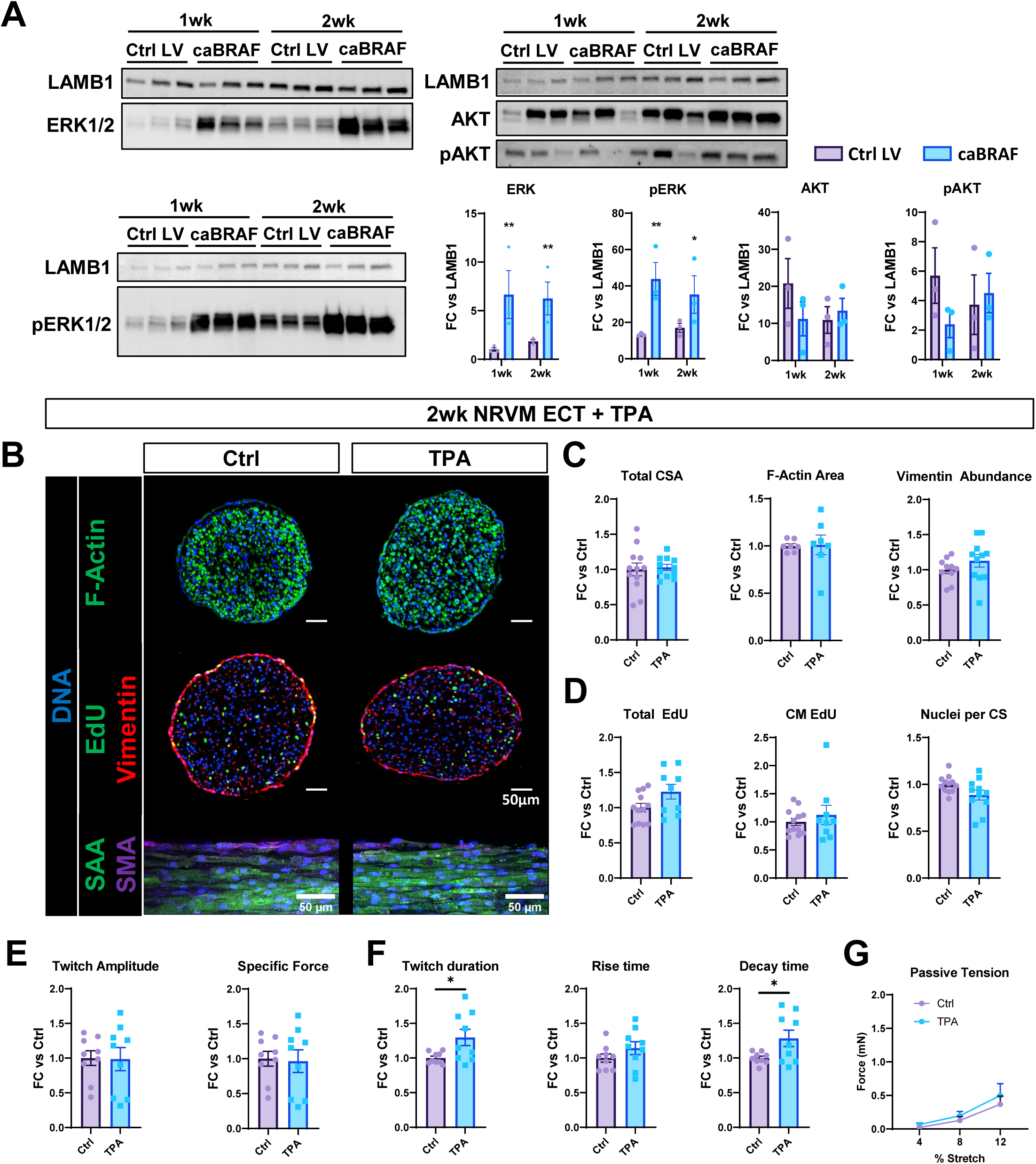
ERK Activation via PKC Stimulation does not Recapitulate caBraf Phenotype in NRVM ECTs. (A) Representative Western blots and corresponding quantifications of ERK and AKT activity in 2 wk ECTs shown normalized to LAMB1. (B-D) Representative cross-section and whole-mount images (B) and corresponding quantifications of morphological (C) and cell cycling (D) parameters in 2 wk ECTs treated on culture days 2-14 with a vehicle solution (Ctrl) or 10 nM TPA. SAA, sarcomeric α-actinin; SMA, smooth muscle actin; CSA, cross-sectional area. (E-G) Quantifications of twitch amplitudes and specific forces (E), twitch kinetic parameters (F), and passive tension-length relationships (G) in 2 wk caBRAF ECTs relative to Ctrl ECTs. %Stretch values are shown relative to the initial testing length. Data: n=3 (A), n=9-12 (C-D), n=9 (E-G). Column graphs showing individual data points, mean ± SEM. Line plots, mean + SEM. *p < 0.05, **p < 0.01, ***p < 0.001 vs. Ctrl LV.

### 3.6 Small molecule inhibition of MEK or ERK decreases cell cycling but does not prevent other phenotypic changes in NRVM ECTs induced by caBRAF expression

We finally explored if caBRAF-induced phenotype in NRVM ECTs could be prevented by MEK or ERK inhibition. Similar to our previous studies with caERBB2 expression^29^, we applied 100nM of the small molecule MEK inhibitor PD0325901 (Meki) or ERK inhibitor SCH772984 (Erki) during the second week of ECT culture during which functional deficit in caBRAF ECTs developed (Fig. 1). Compared to vehicle control, neither Erki nor Meki prevented morphological changes induced by caBRAF expression (Fig. 5A,B, Fig. 5-S1), while both Meki and Erki reduced EdU incorporation in not only caBRAF (Fig. 5C) but also Ctrl (Fig. 5-S2) ECTs, suggesting that ERK activity supports cell cycling in ECTs with or without caBRAF expression. Interestingly, for the dose tested, Meki had stronger negative effect on EdU incorporation than Erki. Consistent with the inability to prevent caBRAF-induced morphological remodeling in ECTs, Meki did not alter contractile strength of caBRAF ECTs, while Erki showed a modest (∼1.9-fold), increase in specific force amplitude compared to vehicle control (Fig. 5D). Neither of the inhibitors altered the twitch kinetics in caBRAF tissues (Fig. 5E). Curiously, Meki also showed a trend towards stiffening of caBraf (Fig. 5F) but not Ctrl (Fig. 5S2) ECTs. Overall, unlike in the case of caERBB2 expression^29^, morphological and functional remodeling following sustained caBRAF expression was only moderately and variably affected by Meki or Erki.

**Figure 5.**
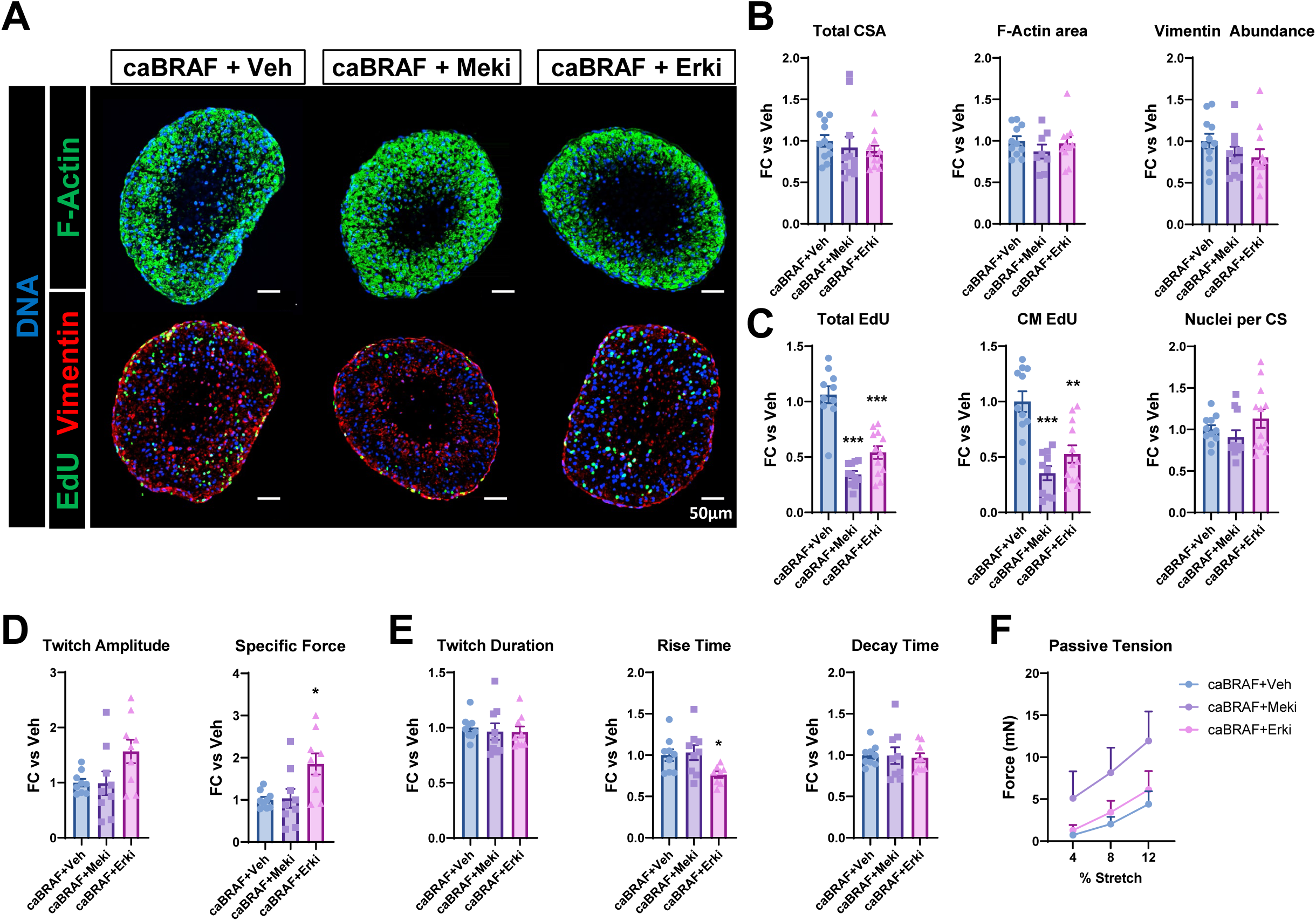
MEK or ERK Inhibition in CaBRAF NRVM ECTs Decreases Cell Cycling without Preventing Morphological or Functional Changes induced by CaBRAF Expression. (A-C) Representative cross-section images (A) and corresponding quantifications of morphological (B) and cell cycling (C) parameters in 2 wk caBRAF ECTs treated on culture days 8-14 with a vehicle solution (caBRAF+Veh) or 100 nM MEK (caBRAF+Meki) or ERK (caBRAF+Erki) inhibitor. CSA, cross-sectional area. (D-F) Quantifications of inhibitor effects on twitch amplitudes and specific forces (D), twitch kinetics parameters (E), and passive tension-length relationships (F) in 2 wk caBRAF ECTs shown relative to caBRAF+Veh group. %Stretch values are shown relative to the initial testing length. Data: n=10-11 (B-C), n=9 (D-F). Column graphs showing individual data points, mean ± SEM. Line plots, mean + SEM. *p < 0.05, **p < 0.01, ***p < 0.001 vs. caBRAF+Veh.

## 4. Discussion

The MAPK pathway is a complex signal transduction pathway with broad effects on cellular fate^52, 54^ and function^18, 55^. In this study, we sought to determine the molecular and functional effects of targeted ERK activation in neonatal rat cardiomyocytes in the context of an *in vitro* engineered heart tissue model. MAPK pathway and ERK activity have been recently implicated in studies attempting to restore heart function after myocardial infarction via endogenous CM proliferation. Thus far, these studies involved activation of MAPK upstream signaling components such as cell receptors (caERBB2)^21, 56^ or their ligands (EGF, FGF, NRG1)^57-60^, but not the direct modulation of downstream components (caBRAF, caERK)^46^. Stimulation of upstream components of the pathway can activate not only the canonical RAF/RAS/MEK/ERK signaling axis, but also parallel mitogenic pathways such as the PI3K/AKT/mTOR^61, 62^. To examine direct ERK-mediated effects on CMs, we expressed BRAF-V600E in NRVMs to constitutively drive ERK activity. As a result, we observed cell-autonomously-induced broad transcriptomic changes leading to sustained CM cycle activity, a shift toward glycolytic metabolism, deteriorated sarcomeres, contractile deficit, and tissue stiffening. In contrast, with time of culture control NRVM tissues underwent gradual functional maturation and exit from the cell cycle. Collectively, our studies suggest that sustained ERK activity induced via BRAF mutagenesis can induce a pro-growth, immature phenotype in neonatal CMs, which warrants further studies in the contexts of congenital heart disease and cardiac regeneration.

One curious finding of this study was that the small molecule PKC activator TPA was unable to recapitulate the morphological and functional changes induced by lentiviral caBRAF expression in NRVM ECTs (Fig. 4B-G), despite increased transcription of ERK-related genes to a similar level seen with caBRAF (Fig. 4-S1B). ERK target genes are known to act within negative feedback loops that inhibit upstream MAPK activity^11, 63, 64^. We observed increased expression of the Sprouty phosphatase *Spry4* following TPA administration, which may attenuate the PKC activation of RAS^65^ or generally reduce MAPK signal flux via inhibition of RAS-GTP formation^66, 67^. Additionally, TPA induced significantly higher expression of the ERK phosphatase *Dusp6*, which could also contribute to the overall attenuation of ERK activity^68^. Similarly, we were surprised to find that caBRAF tissues were largely insensitive to MEK or ERK inhibition (Fig. 5), which we had found previously to be effective in preventing caERBB2-induced contractile deficit and tissue stiffening in NRVM ECTs at the same dosage^29^. This suggests that for caBRAF, higher doses of MEKi or ERKi than expected are required to completely inhibit increased abundance of p-ERK.

Our immunostaining and gene expression analyses were suggestive of both altered ECM composition and cytoskeletal changes contributing to caBRAF tissue remodeling and increased stiffness, but these observations warrant additional studies. Interestingly, in the context of cancer progression^69^, a stiffer extracellular environment induced by RAS-RTK oncogene-expressing cells was necessary to amplify mechanotransduction-induced, YAP/TAZ-dependent tumorigenic cell reprogramming and proliferative growth. A parallel can be drawn between these studies and our observations that by 1wk of culture, the stiffness of caBRAF ECTs was increased while CM cycling remained unchanged, and that by 2 wks of culture and beyond, continued tissue stiffening was accompanied by increased cell cycling (Fig. 1-S4). While modest cell and tissue stiffening are characteristic of natural ECT maturation *in vitro* and postnatal cardiac development *in vivo*^*70*^, they are associated with progressive cell cycle exit rather than sustained CM proliferation. Thus, the temporal dynamics of caBRAF-induced ERK signaling during 4-wk NRVM ECT culture may be qualitatively different from the ERK dynamics during postnatal heart development *in vivo*^*46, 47, 71*^ and will require further mechanistic studies.

In addition to increased ECT stiffness, we consistently found that caBRAF expression drove slower contraction kinetics in a primarily CM-autonomous manner (Fig. 1, Fig. 1-S2,S4), with paracrine factors being a minor contributor to this phenotype (Fig. 3G). In CMs, twitch rise and decay time are significantly influenced by the rate of calcium release from the sarcoplasmic reticulum (SR) via RYR2 receptors and uptake into the SR via the SERCA pump^72^. Both RYR2 and SERCA (ATP2A2) were downregulated in caBRAF ECTs, among other calcium handling genes (Fig. 2-S1), which could contribute to weaker and slower contraction. Additionally, it is possible that sarcomere loss contributed not only to a decrease in twitch amplitude but also to slowed twitch kinetics^73^, although a moderate increase in twitch decay time could be induced by ERK activation in the absence of sarcomere disorganization (Fig. 3 and 4).

Considering ERK is commonly implicated in cardiac hypertrophy^6, 7, 47^, we were intrigued not to observe significantly enlarged size of caBRAF-expressing CMs in flow cytometry analysis of 2D cultures (Fig. 2E) or a prominent hypertrophic signature in RNAseq analysis of 3D ECTs. Instead, the larger size of caBRAF ECTs appeared to be induced by increased numbers of cells, especially at the later timepoints in culture. The CM growth within ECTs may have been in part limited by the lack of vasculature and reliance on the diffusion for oxygen and nutrient supply, which likely led to the development of a central necrotic core^29^ observed in caBRAF ECTs (Fig. 1B,E and Fig. 1-S2A,D). Additionally, despite a difference in stiffness, both 2D and 3D culture environments led to significant sarcomere disassembly in caBraf-expressing CMs. While the relatively soft ECTs could attenuate endogenous ERK signaling at the level of membrane receptors^74^, the constitutively active BRAF-driven intracellular signaling transmission and ERK activation were likely stiffness- and environment-independent. Sarcomere disassembly was also observed in mouse hearts *in vivo* upon transient activation of caERBB2 and ERK in CMs^21, 56^; however, this phenotype was associated with CM hypertrophy as well as AKT co-activation, which was absent in caBRAF ECTs (Fig. 4A). It remains to be studied whether the complex interplay between MAPK/ERK and PI3K/AKT pathway^62, 75, 76^ is differently regulated at different stages of CM maturation and by different methods of MAPK manipulation.

While a metabolic switch from glycolysis to oxidative phosphorylation is a hallmark of CM maturation^77, 78^, reversion back to glycolysis has been suggested as an important step in inducing CM proliferation and heart regeneration *in vivo*^48, 79, 80^. Notably, in tumorigenesis, ERK activation has been shown to promote metabolic reprogramming favoring glycolysis as a means to outcompete neighboring cells for energy^81^. ERK activation via caBRAF expression in NRVM ECTs mirrored that process, as the majority of significantly downregulated GO terms indicated disrupted aerobic metabolism (Fig. 2D), while follow-up studies of ECT-conditioned media confirmed near-exhaustion of glucose and significant lactate accumulation (Fig. 2G). Together, similarly to studies in mouse and zebrafish hearts^48^, ERK activation in NRVMs within ECTs stimulated glycolysis supportive of CM proliferation and tissue growth.

In summary, we have shown that sustained BRAF-V600E-mediated ERK activation in neonatal rat CMs stimulates their glycolytic metabolism, proliferative and migratory capacity, structural remodeling, tissue growth, and contractile deficit. Methods to precisely and temporally manipulate ERK signaling in CMs may represent a promising strategy for therapeutic cardiac repair.

## Data Availability

RNAseq datasets will be made available on GEO at the time of publication, and other data is available upon request.

## Acknowledgements

We thank J. Ou and A. Helfer for consultations regarding our RNA-sequencing analysis, and M. DeLuca for consultations regarding our cell migration analysis.

